# Light Guided *In-vivo* Activation of Innate Immune Cells with Photocaged TLR 2/6 Agonist

**DOI:** 10.1101/128942

**Authors:** Keun Ah Ryu, Bethany McGonnigal, Troy Moore, Rock J. Mancini, Aaron P. Esser-Kahn

**Author notes:** **Present Addresses** Department of Chemistry, University of California, Irvine, Irvine, CA 92697.

## Abstract

The complexity of the immune system creates challenges in exploring its importance and robustness. To date, there have been few techniques developed to manipulate individual components of the immune system in an in vivo environment. Here we show a light-based dendritic cell (DC) activation allowing spatial and temporal control of immune activation in vivo. Additionally, we show time dependent changes in RNA profiles of the draining lymph node, suggesting a change in cell profile following DC migration and indicating that the cells migrating have been activated towards antigen presentation.

## Introduction

Harnessing the innate and adaptive immune response has led to the development of vaccines and therapeutics.^1-3^ However, as the immune system “rivals the nervous system in complexity,^4^” understanding how to design better responses and therapies remains a challenge. One area of complexity is the presentation of antigens by the innate system to the adaptive system – including chemical signaling, spatial migration and cell-cell signaling. During this process, dendritic cells, activated by Toll-like receptors (TLRs) convey pathogenic information to the cells of the adaptive immune systems through the production of cytokines and cell surface markers.^5, 6^ This process involves the migration of activated DCs into lymphatics to present antigens to T-cells.^7-10^ However, understanding this complex system by manipulating sets of cells within it has been a challenge. Chemical control of various innate and adaptive immune cellular processes has been a burgeoning area of interest.^11-15^ Recently, we developed a method to tag and remotely induce a guided immune response (TRIGIR) with a photo-caged TLR2/6 agonist.^16^ TRIGIR allows for selective labeling of cells, followed by remote light activation. Here we use the TRIGIR method for *in vivo* light-based activation to control the migration of dendritic cells. We validate our in vivo activation by monitoring DC migration using adoptively transferred bioluminescent DCs (Luc-DCs) that bear the TRIGIR compound.

Further, to confirm that the migrating cells were presenting antigens and further priming adaptive immune cells^17, 18^, we performed RNA analysis on the target lymph-node. Reported herein is a general procedure where adoptively transferred immune cells can be remotely activated using a UV light source. Though this methodology calls for a TLR2/6 bearing cell type and has limited tissue penetration of UV light used to activate the cells, it may find use in controlling activation of skin or subcutaneous DCs and for studying effects of inflammation within different spatiotemporal parameters. We expect that improvements in both optogenetic techniques, longer wavelength photo-cages, and light delivery methods will help expand the technique to answer many different immunological questions.

## Results

### a. Cell labeling with NPPOC-Pam_2_CSK_4_

Previous work from our lab showed that photo-caging of the N-terminus of the TLR2/6 agonist, Pam_2_CSK_4_^19^, can inhibit its activity to activate TLR2/6. Upon light exposure and subsequent uncaging of the N-terminus, TLR2/6 is activated by the TRIGIR compound. The intercalation of the TRIGIR compound’s palmityl chains^20^ on the TLR2 of dendritic cells allows labelling of the agonists to quiescent innate immune cells without activating TLR2/6. These labelled cells can then be used in adoptive transfer experiments to achieve remote control of inflammatory processes via TLR2.

We sought to adapt this technique in vivo by labeling cells, performing subcutaneous injection and then activating of the cells in their local environment. As the agonist stays co-localized, we can have the spatial control of agonist presentation and immune cell activation.^16^

In initial experiments, we observed that high concentration of the TRIGIR compound, NPPOC-Pam_2_CSK_4_ (**1**, Figure 1a), incubation overnight resulted in higher amount of labeling of the agonist (Figure 1d). However, this also resulted in higher background activation of the cells (Figure 1e). Therefore, labeling the Luc-DCs at 0.1 μM (Figure 1c) showed both good labeling and did not elicit a background immune response. (Figure 1d, e)

**Figure 1.**
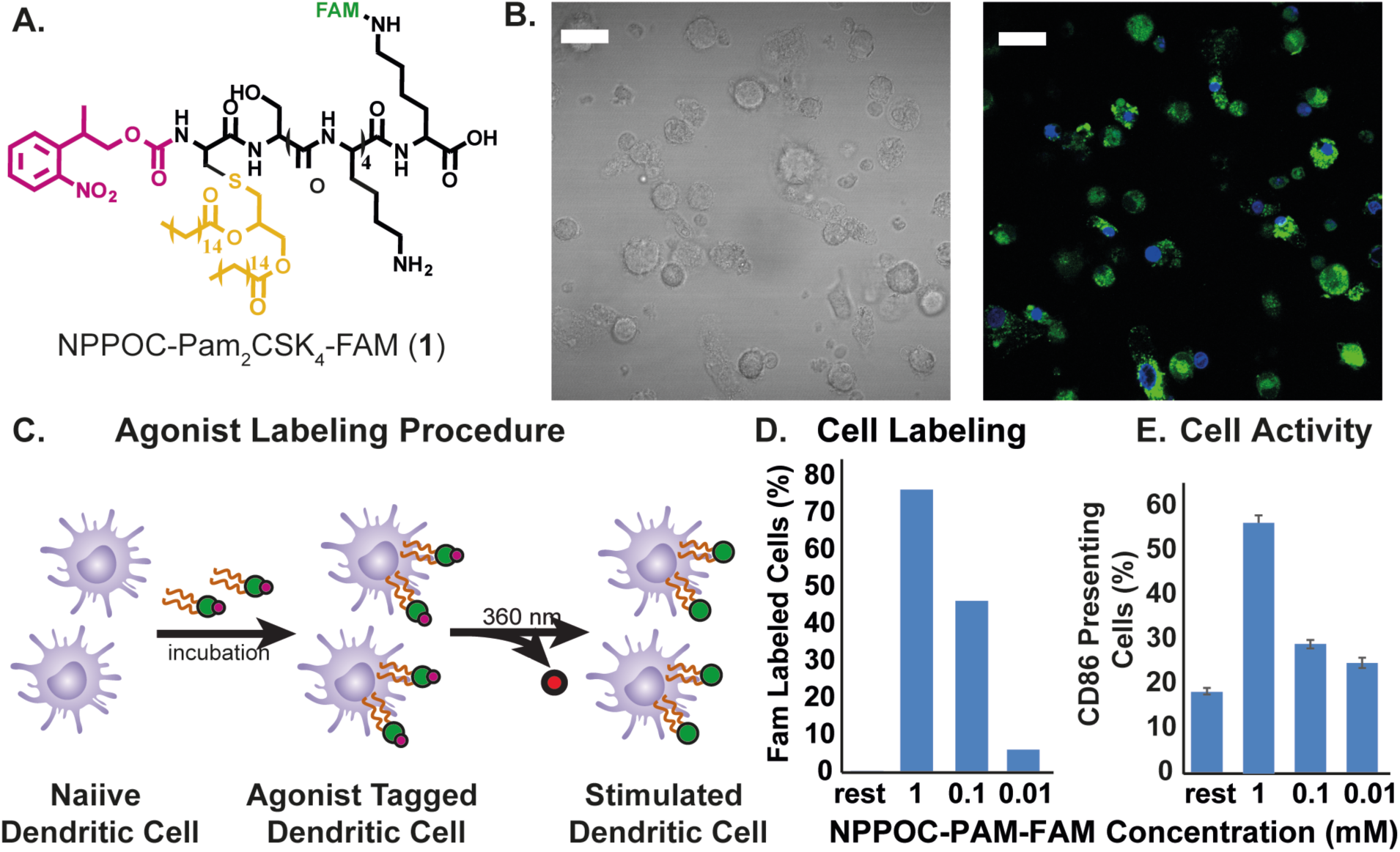
A) Structure of photo-caged TLR 2/6 agonist NPPOC-Pam_2_CSK_4_ (**1**) with fluorescein tag, B) bright field and fluorescent microscopic image of labeled DCs (green-**1**, blue-dapi, scale bar 10 μm), C) NPPOC-Pam-FAM labeling procedure D) Efficiency of agonist labeling, e) background CD86 presentation induced by labeling DCs.

### b. Photo-activation of transferred dendritic cells

Before adoptive transfer, the DCs were incubated with **1** over-night. The cells were then washed to remove excess **1** in the supernatant. The labeled cells were then injected into the footpad of mouse at 1 million cells/30 μL for the mice. To activate the cells with light, the injected footpad of mice was then irradiated with 360 nm light for 15 mins. To determine the limit of activity due to the limit of UV light tissue penetration, we irradiated labelled cells with 360 nm light for 15 min in vitro before injection. This experiment served as a “pre-activated” control and served as an upper limit for what might be achieved with photo-activated DCs in vivo. During the imaging process, following previously reported procedures,^21^ we blocked the bioluminescence occurring from the injected foot with black tape to enhance the signal from the popliteal lymph node. (Figure 2)

**Figure 2.**
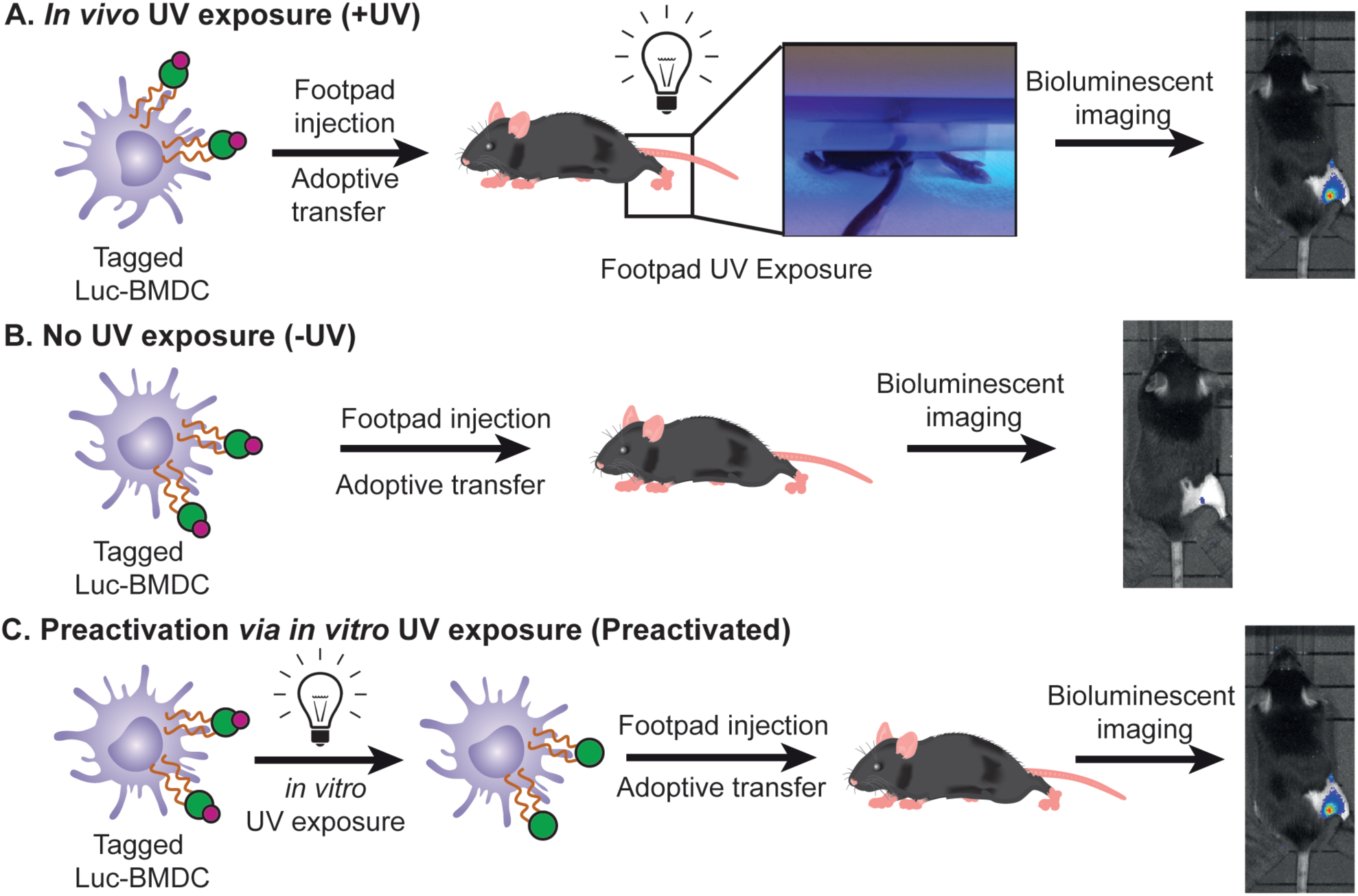
TRIGIR DC adoptive transfer procedures for A) footpad UV irradiated mouse following adoptive transfer, B) mouse with no UV irradiation, and C) Mice injected with pre-irradiated tagged DCs.

To understand the activity of mature DCs, we compared the migration of the Pam_2_CSK_4_ stimulated DCs and non-stimulated DCs that were adoptively transferred into the footpad of a mouse over a period of 96 hrs. We found the Pam_2_CSK_4_ stimulated Luc-DCs migrate faster than the unstimulated Luc-DCs, where we observed migration activity as early as 24 h in Pam_2_CSK_4_ stimulated Luc-DCs with a slow migration, over 96 hrs, of the unstimulated DCs into the draining lymph node at later time points. (Figure 3A, B) Because activation of dendritic cells leads to upregulation of cell surface receptors that aid in the migration and translocation of DCs into the lymph node^22^, we theorize that a shorter time is required for the activated cell to migrate into the lymph node compared to the unstimulated DCs.

**Figure 3.**
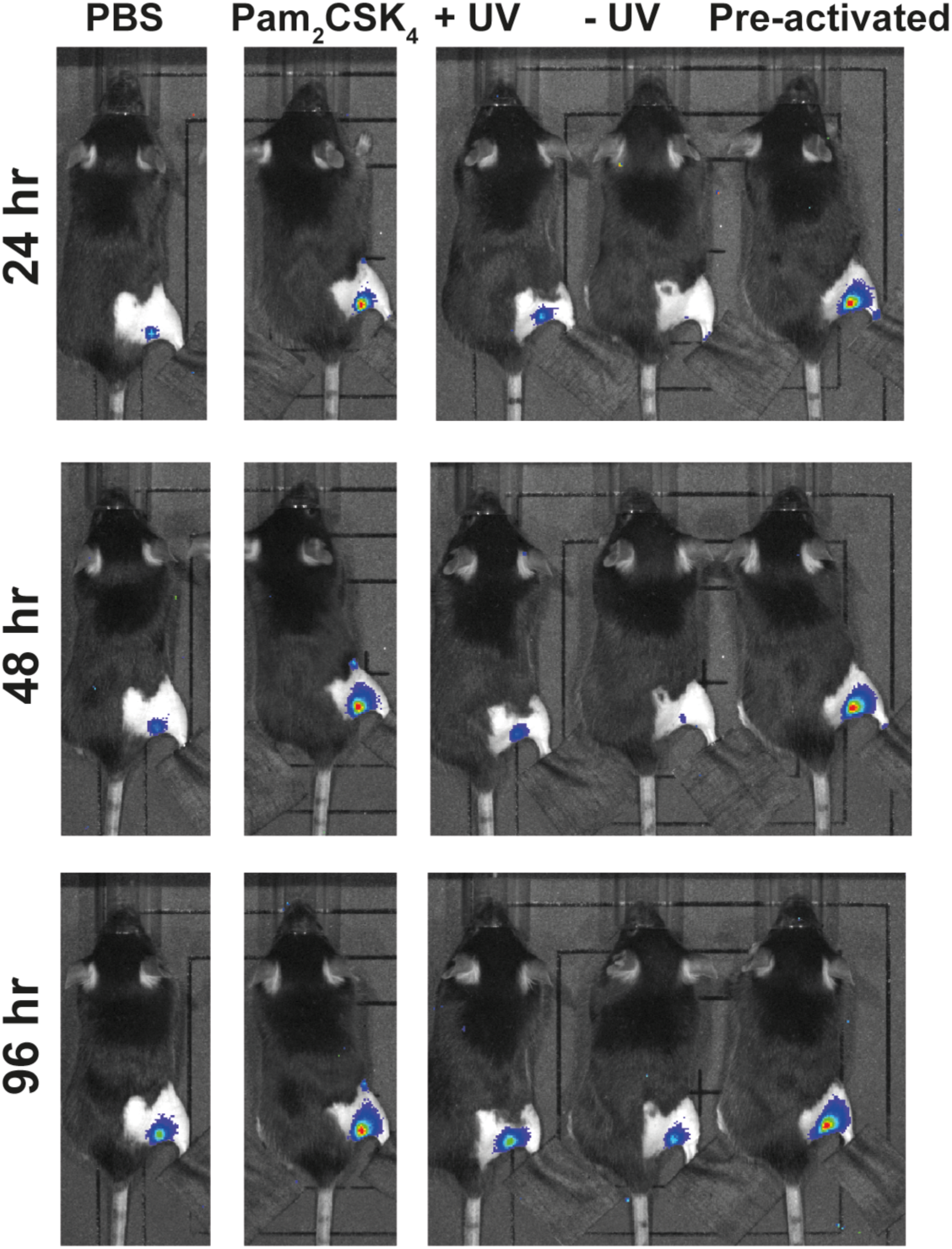
Bioluminescent image of mice taken every 24 h, over 96 h. (A) Control mice include a set of 6 mice injected with DCs preconditioned with Pam2CSK4, and a set of 6 mice injected with DCs with no preconditioning. The test set includes a set of 6 mice with TRIGIR labeled DCs followed by light exposure (+ UV), one with no light exposure (-UV), and a mouse injected with TRIGIR labeled DCs exposed to light before footpad injection (pre-activated).

We sought to determine if light-activation of TLR2 via **1** in vivo recapitulated the migration of activated DCs. We imaged the migration of the Luc-DC in mice whose footpads were exposed to UV light (+UV) or not exposed to UV (-UV). Following the trends seen in the Pam_2_CSK_4_ stimulated cells, the footpads which were directly exposed to UV showed migration of Luc-DCs into the popliteal lymph node much sooner than that of the non-exposed footpads. (Figure 3A, C) Additionally, the cells that were exposed to UV migrate at a similar rate as the cells that were photo-activated before being transferred into a mouse. From this data, we conclude that TRIGIR labelled cells can be activated with light in a non-invasive manner and recapitulate the timing and quantity of their migration to the lymph node.

### c. Confirmation of systemic activation via RNA analysis of popliteal lymph node.

To further confirm the inflammatory state of TRIGIR activated DCs in vivo by light, we harvested popliteal lymph nodes from the mice and analyzed the RNA levels. This measurement also helped us determine if the activated DCs were enacting their antigen presenting role. If the cells were activated following light exposure, the migrated cells will elicit a systemic response as recruitment and maturation of adaptive immune cells occurs in the lymph node. We harvested lymph nodes from both light irradiated and non-irradiated animals which all contained TRIGIR-labeled DCs identical to our previous experiments. To determine differences, we plotted the changes as a relative fold-change of the from irradiated:non-irradiated at each time point. Using this measurement, we determined how irradiation and TLR stimulation changed activity in the lymph node.

**Figure 4.**
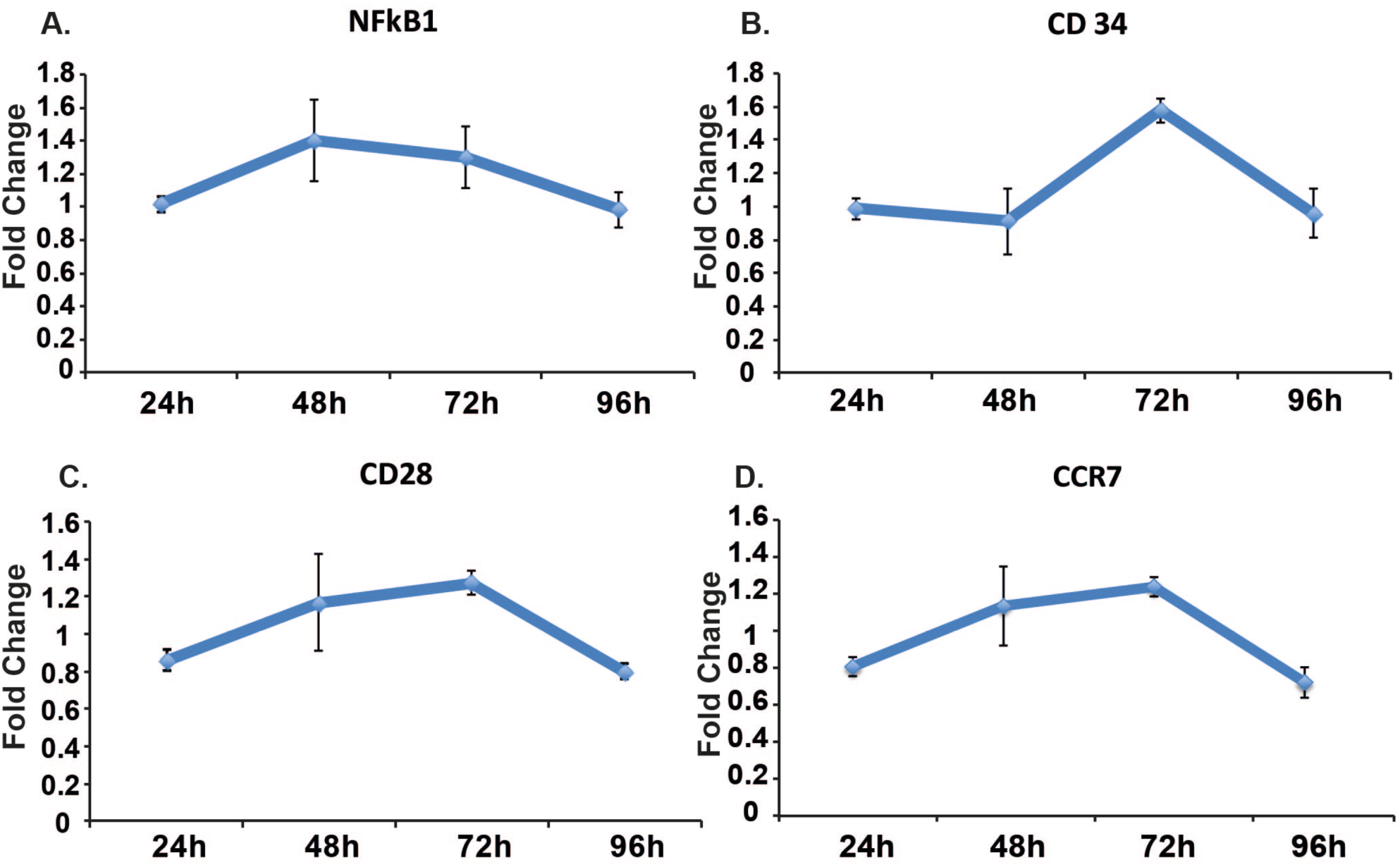
Change in gene profile in harvested lymph node of tested mice. Fold change determined by the ratio of UV irradiated and non-irradiated mice at each time point (n=6) of nfkb1 (a), cd34 (b), cd28 (c), and ccr7 (d).

First, we observed that upon TRIGIR activation, there is a gradual increase in ccr7 which is upregulated by immune cells that enter the lymph node through recognition of CCL19 and CCL21 on the lymph node.^23-25^ From this we conclude there are more ccr7 producing cells recruited into the lymph node. These cells are likely the TRIGIR activated dendritic cells which we observed migrate to the lymph node as well as T cells that have been recruited into the lymph node within the first 72 h after UV exposure as a result of DC activation.

We saw further evidence for T cell recruitment upon TRIGIR activation with an increase of cd34 and cd28 within the same time period. CD34 is required for T cells to enter the lymph node while blocking DC migration into the lymph node.^26^ The downregulation of cd34 at early time points matches the increased migration of the stimulated DCs from the footpad into the lymph node. The gradual increase suggests the increase of T cell trafficking into the lymph node and decrease in DC migration from the footpad.

Similar to the cd34 and cd28 trends, we saw a gradual increase in cd28, a T cell receptor that recognizes CD80 and CD86.^27^, reaching a maximum after 72hrs. This gradual rise indicates the increase in T cell population in the popliteal lymph node. These trends follow known T cell maturation and migration following mature DC contact in the lymph node.^28^

In comparison, there is a general upregulation of nfkb1^29, 30^ starting as early as 48 hours, which could be due to the inflammatory signaling from the activated dendritic cells that have migrated into the popliteal lymph node.

## Discussion

With our method of in vivo photo-activation of immune cells, we delivered a photo-caged, TRIGIR agonist and activated it in a non-invasive manner with light. Using the TRIGIR method of tagging cells, we can overcome the limitation of spatial control of soluble agonists as well as site-specific cell delivery. Compared to conventional adoptive transfer methods that require activation of cells prior to transfer to the animal our method allows for less steps in preparation of the transferred cells and controls when the cells will be activated following adoptive transfer. In addition to temporal control of cell activation, this method offers for the potential of light dosage dependent mitigation of inflammatory signals where longer irradiation times would activate more cells, allowing for sustained activation without the increasing inflammatory response.

This method can also be applied to a variety of cells to induce different responses to TLR2/6 activation. Because TRIGIR is cell specific, but requires labeling, it is compatible with many different primary cell types that can be adoptively transferred. By changing the types of cells and cell populations, one can dissect not only endocrine signaling, but also paracrine signaling following light activation of cell subsets. The technique will not limit researchers to adoptive transfer in the footpad but can create a depot of tagged, subcutaneous cells placed close to an area of interest and gain spatial and temporal control of elicited cellular response. We offer the clear caveat that current photo-activation methods will limit this method to dermal or subcutaneous activation of innate immune cells. Our data suggest that this technique will give researchers the potential to customize an innate cellular response depending on the target disease or immunological model. In conclusion, we present a method for light activation of adoptively transferred cells via TLR2/6. This technique presents a unique way to answer spatial and temporal questions about the innate immune response.

## Associated Content

Supporting Information

Synthesis of photocaged agonist, biological protocols, additional cell assay data are available in the supporting information.

## AUTHOR INFORMATION

### Author Contributions

K.R. designed and performed experiments, analyzed data, and wrote the manuscript. R.J.M. initially synthesized the photocaged agonist. B. M. and T. M. performed mouse experiements.

A.P.E. supervised the project. All authors provided comments and contributions and have given approval to the final version of the manuscript.

### Notes

The authors declare no competing financial interests.

## Achnowledgement

We would like to thank the Prescher Laboratory for help with IVIS imaging.

## Reference

1. Querec, T.; Bennouna, S.; Alkan, S. K.; Laouar, Y.; Gorden, K.; Flavell, R.; Akira, S.; Ahmed, R.; Pulendran, B., Yellow fever vaccine YF-17D activates multiple dendritic cell subsets via TLR2, 7, 8, and 9 to stimulate polyvalent immunity. J. Exp.Med. 2006, 203 (2), 413–424.

2. Schreibelt, G.; Tel, J.; Sliepen, K.; Benitez-Ribas, D.; Figdor, C.; Adema, G.; de Vries, I., Toll-like receptor expression and function in human dendrtiic cell subsets:implications for dendritic cell-based anti-cancer immunotherapy. Cancer Immunol. Immunother. 2010, 59, 1573–1582.

3. Melero, I.; Berman, D.; Aznar, A.; Korman, A.; Gracia, J.; Haanen, J., Evolving synergistic combinations of targeted immunotherapies to combat cancer. Nat. Rev. Cancer 2015, 15, 457–472.

4. Adler; M., E., Signaling Breakthroughs of the Year. Sci. Signal. 2017, 10, eaam5681.

5. Katsikis, P.; Schoenberger, S.; B., P., Probing the 'labyrinth' linking the innate and adaptive immune systems. Nat. Immunol. 2007, 8, 899–901.

6. Iwasaki, A.; Medzhitov, R., Toll-like receptor control of the adaptive immune responses. Nat Immunol 2004, 5 (10), 987–995.

7. Schimmelpfennig, C. H.; Schulz, S.; Arber, C.; Baker, J.; Tarner, I.; McBride, J.; Contag, C. H.; Negrin, R. S., Ex Vivo Expanded Dendritic Cells Home to T-Cell Zones of Lymphoid Organs and Survive in Vivo after Allogeneic Bone Marrow Transplantation. Am. J. Pathol. 2005, 167 (5), 1321–1331.

8. Randolph, G. J.; Angeli, V.; Swartz, M. A., Dendritic-cell trafficking to lymph nodes through lymphatic vessels. Nat. Revi. Immunol. 2005, 5 (8), 617–628.

9. Wu, W.; Li, R.; Malladi, S. S.; Warshakoon, H. J.; Kimbrell, M. R.; Amolins, M. W.; Ukani, R.; Datta, A.; David, S. A., Structure-Activity Relationships in Toll-like Receptor-2 agonistic Diacylthioglycerol Lipopeptides. J. Med. Chem. 2010, 53 (8), 3198–3213.

10. Buwitt-Beckmann, U.; Heine, H.; Wiesmüller, K.-H.; Jung, G.; Brock, R.; Ulmer, A. J., Lipopeptide structure determines TLR2 dependent cell activation level. FEBS J. 2005, 272 (24), 6354–6364.

11. Nico J. Meeuwenoord;. Prof. Ferry A. Ossendorp; Prof. Herman S. Overkleeft; Dr. Dmitri V. Filippov; Dr. Sander I. van Kasteren, Pawlak, J.; Geoffroy, G.; Ruckwardt, T.; Bremmers, J.; Meeuwenoord, N.; Ossendorp, F.; Overkleeft, H.; Filippov, D.; van Kasteren, S., Bioorthogonal Deprotection on the Dendritic Cell Surface for Chemical Control of Antigen Cross-Presentation. 2015, 54, 5628–5631.

12. Parasar, B.; Chang, P. V., Chemical optogenetic modulation of inflammation and immunity. Chem. Sci. 2017, 8, 1450–1453.

13. Liu, H.; Kwong, B.; Irvine, D. J., Membrane Anchored Oligonucleotides for in vivo Tumor Cell Modification and Localized Cancer Immunotherapy. Angew. Chem. Int. Ed. 2011, 50, 7052–7055.

14. Govan, J. M.; Young, D. D.; Lively, M. O.; Deiters, A., Optically Triggered Immune Response through Photocaged Oligonucleotides. Tetrahedron Lett. 2015, 56, 3639–3642.

15. Parker, C. G.; Domaoal, R. A.; Anderson, K. S.; Spiegel, D. A., An Antibody-Recruiting Small Molecule That Targets HIV gp120. J. Am. Chem. Soc. 2009, 131, 16392–16394.

16. Mancini, R. J.; Stutts, L.; Moore, T.; Esser-Kahn, A. P., Controlling the Origins of Inflammation with a Photoactive Lipopeptide Immunopotentiator. Angew. Chemie. Int. Ed. 2015, 54 (20), 5962–5965.

17. Martín-Fontecha, A.; Lanzavecchia, A.; Sallusto, F., Dendritic Cell Migration to Peripheral Lymph Nodes. In Dendritic Cells, Lombardi, G.; Riffo-Vasquez, Y., Eds. Springer Berlin Heidelberg: 2009; pp 31–49.

18. Martín-Fontecha, A.; Sebastiani, S.; Höpken, U. E.; Uguccioni, M.; Lipp, M.; Lanzavecchia, A.; Sallusto, F., Regulation of Dendritic Cell Migration to the Draining Lymph Node. J. Exp. Med. 2003, 198 (4), 615–621.

19. Spohn, R.; Buwitt-Beckmann, U.; Brock, R.; Jung, G.; Ulmer, A. J.; Wiesmüller, K.-H., Synthetic lipopeptide adjuvants and Toll-like receptor 2—structure–activity relationships. Vaccine 2004, 22 (19), 2494–2499.

20. Kang, J. Y.; Nan, X.; Jin, M. S.; Youn, S.-J.; Ryu, Y. H.; Mah, S.; Han, S. H.; Lee, H.; Paik, S.-G.; Lee, J.- O., Recognition of Lipopeptide Patterns by Toll-like Receptor 2-Toll-like Receptor 6 Heterodimer. Immunity 2009, 31 (6), 873–884.

21. Lee, H. W.; Yoon, S. Y.; Singh, T. D.; Choi, Y. J.; Lee, H. J.; Park, J. Y.; Jeong, S. Y.; Lee, S.-W.; Ha, J.- H.; Ahn, B.-C.; Jeon, Y. H.; Lee, J., Tracking of dendritic cell migration into lymph nodes using molecular imaging with sodium iodide symporter and enhanced firefly luciferase genes. Sci. Rep. 2015, 5, 9865.

22. Bertho, N.; Adamski, H.; Toujas, L.; Debove, M.; Davoust, J.; Quillien, V., Efficient migration of dendritic cells toward lymph node chemokines and induction of TH1 responses require maturation stimulus and apoptotic cell interaction. Blood 2005, 106 (5), 1734–1741.

23. Ritter, U.; Wiede, F.; Mielenz, D.; Kiafard, Z.; Zwirner, J.; Körner, H., Analysis of the CCR7 expression on murine bone marrow-derived and spleen dendritic cells. J. Leukoc. Biol. 2004, 76 (2), 472–476.

24. Noor, S.; Habashy, A. S.; Nance, J. P.; Clark, R. T.; Nemati, K.; Carson, M. J.; Wilson, E. H., CCR7-Dependent Immunity during Acute Toxoplasma gondii Infection. Infect. Immun. 2010, 78 (5), 2257–2263.

25. Clatworthy, M. R.; Aronin, C. E. P.; Mathews, R. J.; Morgan, N. Y.; Smith, K. G. C.; Germain, R. N., Immune complexes stimulate CCR7-dependent dendritic cell migration to lymph nodes. Nature Medicine 2014, 20 (12), 1458–1463.

26. Drew, E.; Merzaban, J. S.; Seo, W.; Ziltener, H. J.; McNagny, K. M., CD34 and CD43 Inhibit Mast Cell Adhesion and Are Required for Optimal Mast Cell Reconstitution. Immunity 2005, 22 (1), 43–57.

27. Linterman, M. A.; Denton, A. E.; Divekar, D. P.; Zvetkova, I.; Kane, L.; Ferreira, C.; Veldhoen, M.; Clare, S.; Dougan, G.; Espéli, M.; Smith, K. G. C., CD28 expression is required after T cell priming for helper T cell responses and protective immunity to infection. eLife 2014, 3, e03180.

28. Mempel, T. R.; Henrickson, S. E.; von Andrian, U. H., T-cell priming by dendriticcells in lymph nodes occurs in three distinct phases. Nature 2004, 427 (6970), 154–159.

29. Baltimore, D., Discovering NF-κB. Cold Spring Harb. Perspect. Biol. 2009, 1 (1), a000026.

30. Gerondakis, S.; Siebenlist, U., Roles of the NF-κB Pathway in Lymphocyte Development and Function. d Spring Harb. Perspect. Biol. 2010, 2 (5), a000182.

